# Occurrence and distribution of fecal indicators and pathogenic bacteria in seawater and *Perna perna* mussel in the Gulf of Annaba (Southern Mediterranean)

**DOI:** 10.1101/2020.10.04.325167

**Authors:** Mouna Boufafa, Skander Kadri, Peter Redder, Mourad Bensouilah

**Author notes:** Corresponding authors: Mouna Boufafa; Peter Redder.

## Abstract

The brown mussel *Perna perna* is a marine bivalve that is widely distributed and consumed along the east coast of Algeria. Due to its filter-feeding capacity, this mollusk can accumulate large quantities of pathogenic microorganisms from the surrounding waters, thus acting as bio-indicator of coastal environments. The objective of this study is to investigate the occurrence and distribution of fecal indicators and pathogenic bacteria in seawaters and mussels collected from four different sites in the Gulf of Annaba through physicochemical, biochemical and molecular analysis. The obtained results revealed that the levels of fecal indicator bacteria (FIB) were alarmingly high at Sidi Salem and Rezgui Rachid when compared with the two other sites (p < 0.05) and largely exceeded the permissible limits. Besides, *P. perna* collected from all sites were several fold more contaminated by these germs than seawater samples, notably, during the warm season of the study period. Biochemical and molecular analysis showed that isolated bacteria from both environmental compartments were mostly potentially pathogenic species such as *E. coli*, Salmonella, Staphylococcus, Klebsiella, Pseudomonas and Proteus. These principal findings demonstrate the strong involvement of anthropogenic activities on the microbiological quality of the Gulf and highlight the role of *P. perna* as an effective bio-indicator of the bacteriological quality of coastal waters.

## Introduction

For many decades, the coastal marine ecosystems have been continuously threatened by several anthropogenic activities such as improper sewage disposal, urban runoff and massive discharges of agricultural and industrial effluents (Ghozzi et al. 2017; Damak et al. 2020). Coastal waters are often the receiving environment for all kinds of wastewater discharges containing many microorganisms that are harmful to human health, especially in bathing beaches and shellfish production areas (Perkins et al. 2014). Thus, the impact on health is more than worrying, placing microbiological pollution as a major public health problem.

Due to their sessile life-style, resistance to environmental stressors and efficient filtration ability, bivalves, especially mussels, have been widely used as bio-indicators of coastal pollution (Belabed et al. 2013; Jia et al. 2018; Ozkan et al. 2017). These invertebrates have the potential to accumulate large quantities of microorganisms from their surrounding waters, including opportunistic bacteria (Aeromonas, Vibrio, Pseudomonas), protozoan parasites (Cryptosporidium, Giardia), viruses (adenoviruses, hepatoviruses) as well as pathogenic bacteria (*E. coli*, Salmonella) (Ghozzi et al. 2017). They may therefore jeopardize human health, especially when they are consumed as seafood (Stabili et al. 2005; Zannella et al. 2017, Vincy et al. 2017). Numerous studies have reported that many serious illnesses such as acute gastroenteritis and hepatitis E virus infections are related to the presence of pathogenic microorganisms in bivalves mollusks, especially when they are eaten raw or undercooked (Le Guyader et al. 2006; O’Hara et al. 2018; Kobayashi et al. 2019; Fouillet et al. 2020). Hence, there is an urgent need for an overall assessment to predict the presence of these infectious agents related to waterborne outbreaks, and to prevent the impacts of fecal contamination on human and environmental health. The Gulf of Annaba is one of the most valuable coastal regions of Northern Algeria, because of its great touristic and economic importance. However, it is highly vulnerable to several types of pollutants, primarily related to the intensive agricultural and industrial discharges and the presence of domestic wastes, especially on the outskirts of the city where a high number of population is concentrated (Soltani et al. 2012; Amri et al. 2017; Ouali et al. 2018). These anthropogenic sources are further exacerbated by diverse natural environmental contaminants such us terrestrial effluents especially in rainy weather, animal excreta, freshwater and river discharges, and the problem of global climate change. Despite this increasing pressure, the problem of fecal contamination, and the potential health hazards it can cause have been little studied in the Gulf of Annaba (Kadri et al. 2015, 2017). Therefore, this study aimed (1) to evaluate the occurrence and the distribution of fecal indicators and pathogenic bacteria in seawater and the mussel *Perna perna* samples by implementing a spatial-temporal sampling strategy (2) to assess the impact of physicochemical variables on the abundance of fecal indicator bacteria (FIB) (3) to determine the origin of microorganism’s inputs and hot spots of heavy contamination.

## Materials and methods

### Sampling area

The Gulf of Annaba in Northeastern Algeria, stretching ~40 km from Cap de Garde (36°96’N, 7°79’E) in the West to Cap Rosa (36°68’N, 8°25’E) in the East, is a heavily polluted ecosystem, due to a variety of agricultural, industrial and urban discharges, in addition to massive domestic wastes from a large part of the city of Annaba (Abdennour et al. 2000). Four sampling sites were strategically selected for the present study, based on different potential pollution sources in these areas: S1 ‘Cap de Garde’ (36°96’N, 7°79’E); S2 ‘Rezgui Rachid’ (36°91’N, 7°76’E); S3 ‘Sidi Salem’ (36°86’N, 7°76’E); and S4 ‘Lahnaya’ (36°93’N, 8°20’E) (Fig. 1).

**Fig. 1.**
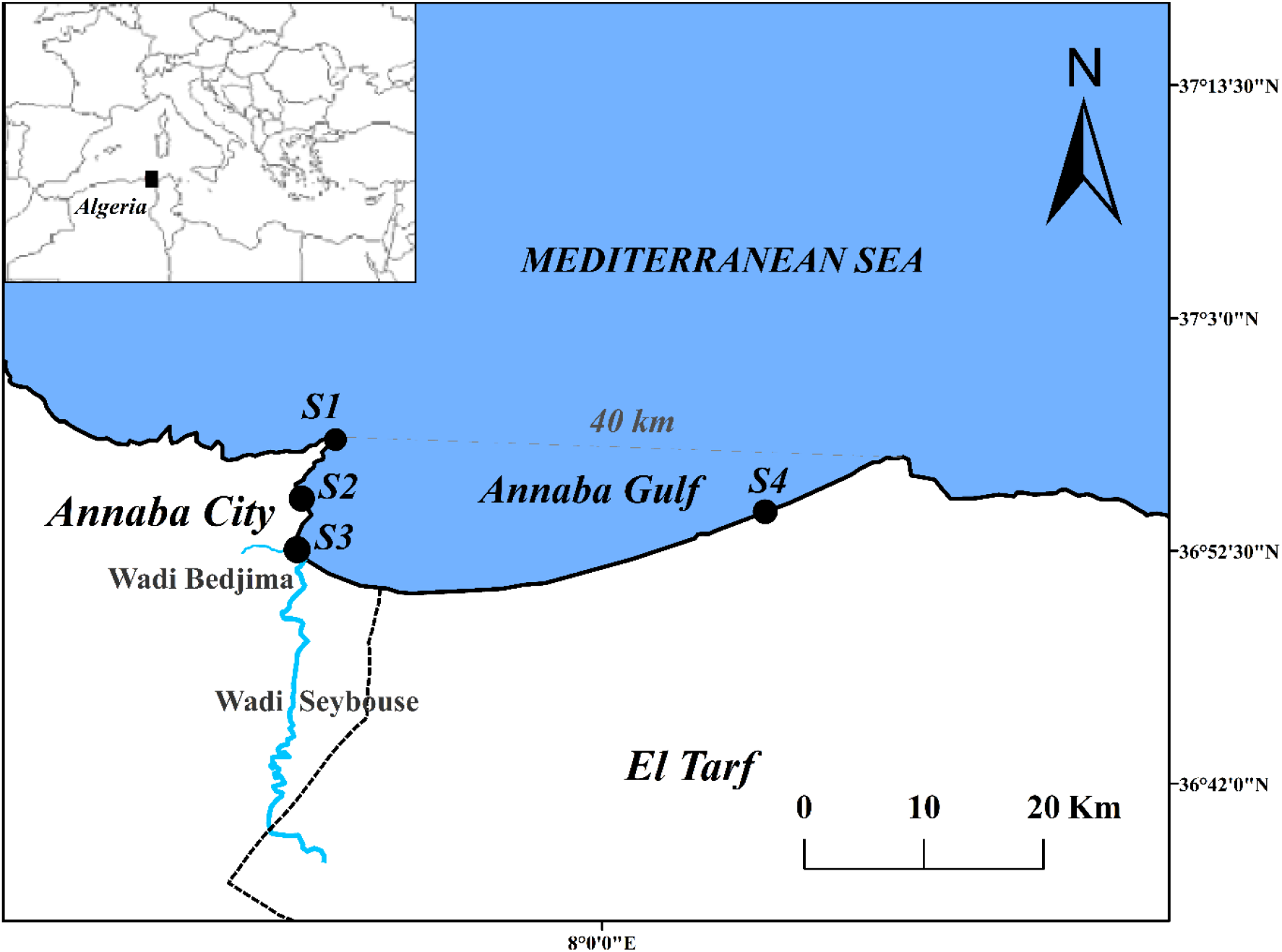
Map showing the location of the Gulf of Annaba and sampling sites. The location of the larger map is shown by a black rectangle on the insert map. The four sampling sites (S1 to S4) are indicated with black circles, S1: Cap de Garde, S2: Rezgui Rachid, S3: Sidi Salem, S4: Lahnaya.

### Sampling protocol

Samples of seawater and *Perna perna* mussels were monthly and simultaneously collected at each site, in the period from January to December 2018. Water samples were obtained at a depth of 30-50 cm below the surface of the water to avoid sunlight exposure using 250 ml sterile glass bottles. *P. perna* mussels were harvested by hand near to the water collecting points at a rate of 10-20 individuals (depending on size). All samples were immediately placed in a clean cooler containing ice cubes (4°C) and transported to the laboratory within the following 2-4h. At each site and each month, seawater environmental variables including temperature (T), pH, salinity (Sal), dissolved oxygen (DO) were measured *in situ* using a multi-parameter probe (Multi 340i/SET-82362, WTW®, Germany). The determination of seawater suspended solids (SS) was performed as described by Aminot and Chaussepied (1983).

### Bacteriological analysis

Once in the laboratory, the mussels were washed and opened aseptically. Then, the tissue and intravalvular liquid (25 g) were mixed and homogenized with 225ml sterile physiological water in a sterile laboratory blender (standard NF EN ISO 6887). For seawater samples, a volume of 100 ml was directly analysed without any treatment (Rodier et al. 2009). The levels of FIB such as total coliforms (TC), *Escherichia coli* (EC) as well as fecal streptococci (FS) were estimated by a three-tube decimal dilution using the most probable number (MPN) method (standard NF V 08-021 (1993) / ISO 7402 and NF V 08-020 (1994) / ISO 7251). All results were statistically expressed as MPN per 100 ml of the sample according to the Mac Grady’s tables (Rodier et al. 2009). As for the isolation of potentially pathogenic bacteria, standard microbial methods were carried out (Rodier et al. 2009). Bacterial isolates were biochemically identified at the species level through Analytical Profile Index (API 20E, API20NE, API Staph) and further confirmed by 16S rRNA gene sequencing, Multi Locus Sequence Typing (MLST), and phylogenetic analysis.

### DNA extraction and 16S rRNA gene amplification

All primers used in this study are listed in Table 1. For cells disruption and DNA extraction from 25 selected isolates, bacterial colonies were picked from pure overnight LB (Luria Bertani) agar plates and transferred into 1.5 ml Eppendorf tubes containing 50 μl of 1xTE buffer (10 mM Tris-HCl pH 8, 1 mM EDTA) supplemented with approximately 100 mg of 0.1 mm Zirconia beads. The tubes were incubated at 37°C for 15 minutes and then strongly vortexed for 3 min to disrupt the cells. The resulting bacterial lysate served as a template for the 16S rRNA gene amplification. The 25 μl PCR mixture contained 0.5 μl DNA template, 2.5 μl Dream Taq buffer (10x), 1.5 μl dNTPs (2.5 mM each), 0.5μl Dream Taq DNA polymerase (Thermo Scientific™) and 1.5 μl 10 μM of each universal primers 27F and 1492R. PCR cycling conditions were maintained as previously described by da Silva et al. (2013). Amplification products were visualized by electrophoresis on 1% agarose gel in 1x TBE buffer after staining with SYBR Safe (Invitrogen) and subsequently purified with Gene Jet Gel Extraction Kit (Thermo Scientific^™^)

### DNA sequence analysis

The PCR-amplified regions of the 16S rRNA genes were Sanger sequenced using primer 27F (Table 1). The obtained partial sequences of the 16S rRNA gene were first analysed and assembled using BioEdit version 7.2.5 (Hall 1999), and then compared with the GenBank NCBI database through BLAST software to confirm the species of the isolates. After that, a multiple sequence alignment was carried out using Clustal X software integrated into MEGA 7 program (Kumar et al. 2016). Finally, the phylogenetic tree was constructed by the neighbor-joining method with 1,000 bootstrap replications.

### Multilocus Sequence Typing analysis (MLST analysis)

Five isolates (EM3, EM18, EM97, EM102, and MM6) isolated from *P. perna*, were chosen based on their same site of isolation (Sidi Salem) and sampling date (15 January 2018) for further characterization by Multi-locus sequence typing (MLST). DNA from all isolates were subjected to PCR amplification targeting seven specific genes (*trpA, trpB, dinB, polB, putP, pabB and icdA*) using suitable primers (Table 1), and following the same procedures as used for 16S rRNA genes. The amplification program was carried out as follow: initial denaturation of 4 min at 94°C, followed by 30 cycles of 30s at 94°C, 30s at 52°C and 2 min at 72°C, and a final extension at 72°C for 4 min. The phylogenetic tree is based on 2758 bp concatenated partial sequences of the seven genes from EM3, EM18, EM97, EM102, and MM6 as well as the equivalent loci in closely related strain. The sequences were aligned with Clustal Omega with default settings on the EBI server and the guide-tree was visualized using iTOL (Letunic and Bork, 2019; Madeira et al. 2019).

### Statistical analysis

Statistical analysis was accomplished under R software version 3.1.2. First, the Spearman correlation coefficient was evaluated to investigate possible relationships between our data sets. Then, the Kruskal-Wallis test was applied to assess the inter-sites and inter-months comparisons. Finally, Principal Component Analysis (PCA) was used as a descriptive method to characterize the structure of the four sampling sites in the study area and to assess the contribution of measured environmental variables on the abundance of the fecal indicators employing the FactoMineR package. In all tests, the significance level was set to p-value < 0.05.

## Results

### Physicochemical analysis of sampled water

The monthly variation in seawater environmental variables obtained throughout the sampling period are presented in Fig. 2. As expected, the annual temperature and salinity cycles showed similar seasonal fluctuations across the four study sites. Seawater temperature ranged from 10.3°C at S4 in February to 28.6°C at S2 in August, while salinity varied from 34.9 g/L at S3 in March to 41.6 g/L at S2 in August. The variations of these two parameters are primarily influenced by the climatic conditions of the area, the high values recorded at S2 and S3 would be due to the fact that these sites, located in the inner of the Gulf, are protected from currents and have low freshwater inputs and fairly high evaporation. The pH remained relatively constant and alkaline during the sampled months, with a slight increase in spring. Changes in dissolved oxygen were generally opposite to the changes in temperature and salinity. The highest value (12.6 mg/L) was recorded during the winter at S4, while the lower one (5 mg/L) was detected during the summer at S3 and S2. Indeed, the application of the Spearman’s correlation test led us to confirm the strong negative and significant correlations between this variable and the temperature and the salinity (r = −0.84, p < 0.0001; r = −0.65, p < 0.0001 respectively) (Table 3). Levels of suspended solids were lower at S1 and S4 as compared with the other two sites. The highest value (0.42mg/L) was recorded two times in February at S3 and in December at S2.

**Fig. 2.**
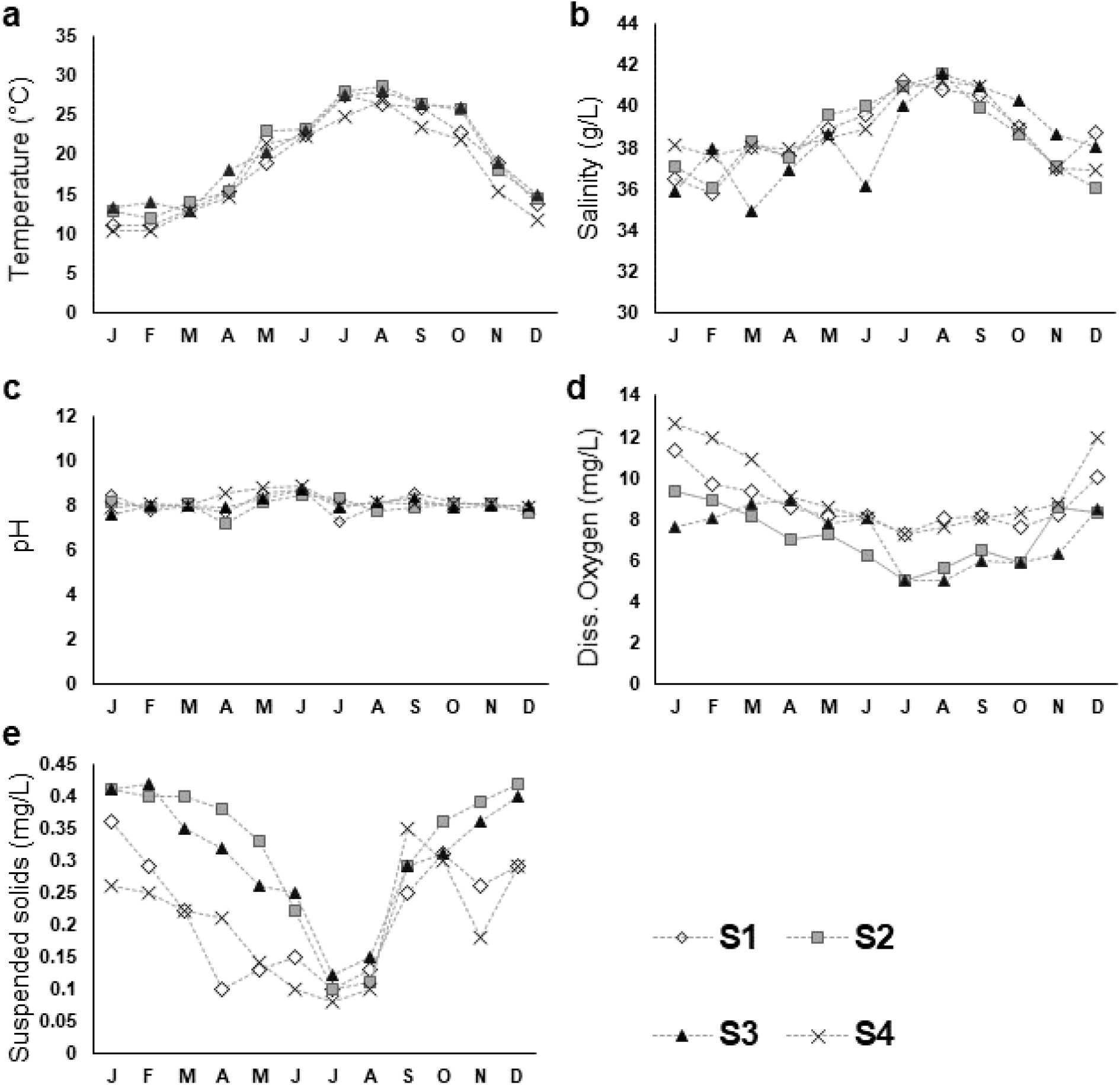
Results of physicochemical analysis of seawater samples at the four sampling sites. **a** Temperature (°C), **b** Salinity (g/L), **c** Dissolved Oxygen (mg/L), **d** pH, **e** Suspended Solids (mg/L). S1: Cap de Garde, S2: Rezgui Rachid, S3: Sidi Salem, S4: Lahnaya.

### Bacteriological analysis of isolated bacteria

As shown in Fig. 3, the results of the bacteriological analysis revealed that the fecal contamination varied over time and between the four sampling sites in the Gulf of Annaba (p < 0.05). However, the levels of FIB were consistently alarmingly high at S2 and S3, and largely exceeded the limits defined by Algerian law. In addition, *P. perna* mussels from all sites were several fold more contaminated by these germs than the seawater samples, notably, during the warm months of the year. TC concentrations in the mussels ranged from 9×10^3^ MPN/100 ml in March at S4 to 3×10^5^ MPN/100mL in June at S3. For seawater samples, the maximum levels of TC (4.6×10^3^ MPN/100 ml) was registered in S2 and S3 where more than 500 MPN/100mL were noted in 96% of the samples. Fecal contamination with *E. coli* was less significant than TC contamination during the sampling period. This may be due to the exclusively fecal origin of this member of the TC group, which makes it probably one of the best bacterial indicators of fecal contamination in the aquatic environment. This germ was present in 44% of the seawater samples at concentrations below 100 MPN/100mL; however, 100% of the *P. perna* samples of S3 showed loads of more than 4.6×10^4^ MPN/100mL. On the other hand, FS was present throughout the entire study period. The highest concentration (2.5×10^4^ MPN/100ml) was detected in September in the mussels of S2. Loads of more than 100 MPN/100mL were noted in more than 79% of the seawater samples of all sites. Based on Spearman’s correlation results, these three groups of bacteria were highly correlated with each other (p < 0.0001).

**Fig. 3.**
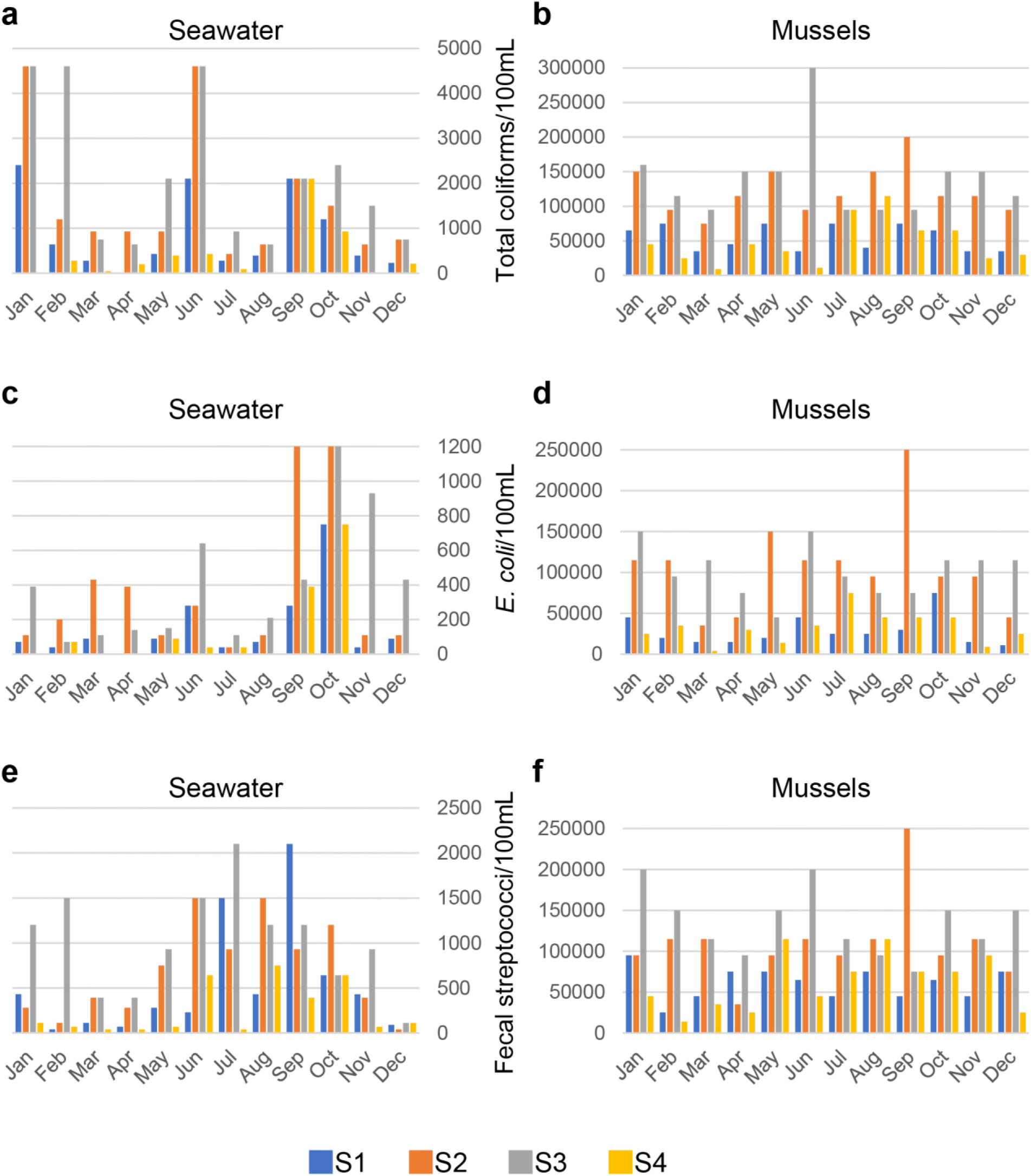
Temporal variations of fecal indicator bacteria in seawater and mussels. **a** and **b** Total coliforms per 100 mL, **c** and **d** *Escherichia coli* per 100 mL, **e** and **f** Fecal streptococci per 100 mL. Note that the scales are different on each diagram. S1: Cap de Garde, S2: Rezgui Rachid, S3: Sidi Salem, S4: Lahnaya.

### Pathogenic bacteria

During the entire study period, a total of 208 bacterial isolates (142 from mussels and 66 from seawater) belonging to 22 genera and 46 species were identified using biochemical tests. The most ubiquitous and abundant microorganism among all the environmental samples was *E. coli* (41.4%), followed by *Aeromonas hydrophila* (5.8%), *Klebsiella pneumoniae* (3.9%), *Pseudomonas aeruginosa* (3.4%), *Enterobacter cloacae* (2.9%), *Vibrio parahaemolyticus* (2.9%), *Burkholderia cepacia* (2.4%), *Morganella morganii* (2.4%), *Micrococcus spp* (1.9%) *Pseudomonas luteola* (1.9%), *Staphylococcus sciuri* (1.9%), *Staphylococcus xylosus* (1.9%), *Providencia rettgeri* (1.4%), *Salmonella spp* (1.4%) and *Yersinia enterolittica* (1.4%) (Only species contributing more than 1% in at least one sample are shown in Fig. 4). In the seawater samples of S1 and especially of S4, the number of pathogenic bacteria did not exceed 14 germs, whereas in S2 and S3, their number was in the order of 16 and 27, respectively. In *P. perna* samples, the number of these infectious agents was 16 in S4, 24 in S1, and 43 and 59 in S2 and S3, respectively. (Table S1). The members of the family *Enterobacteriaceae* were importantly dominant and presented the highest occurrence of all potentially pathogenic microorganisms (65.38%). This large group of bacteria was extensively found in all samples of *P. perna* mussels and its presence was most pronounced in S2 and S3 (Fig. 4; Table S1).

**Fig. 4.**
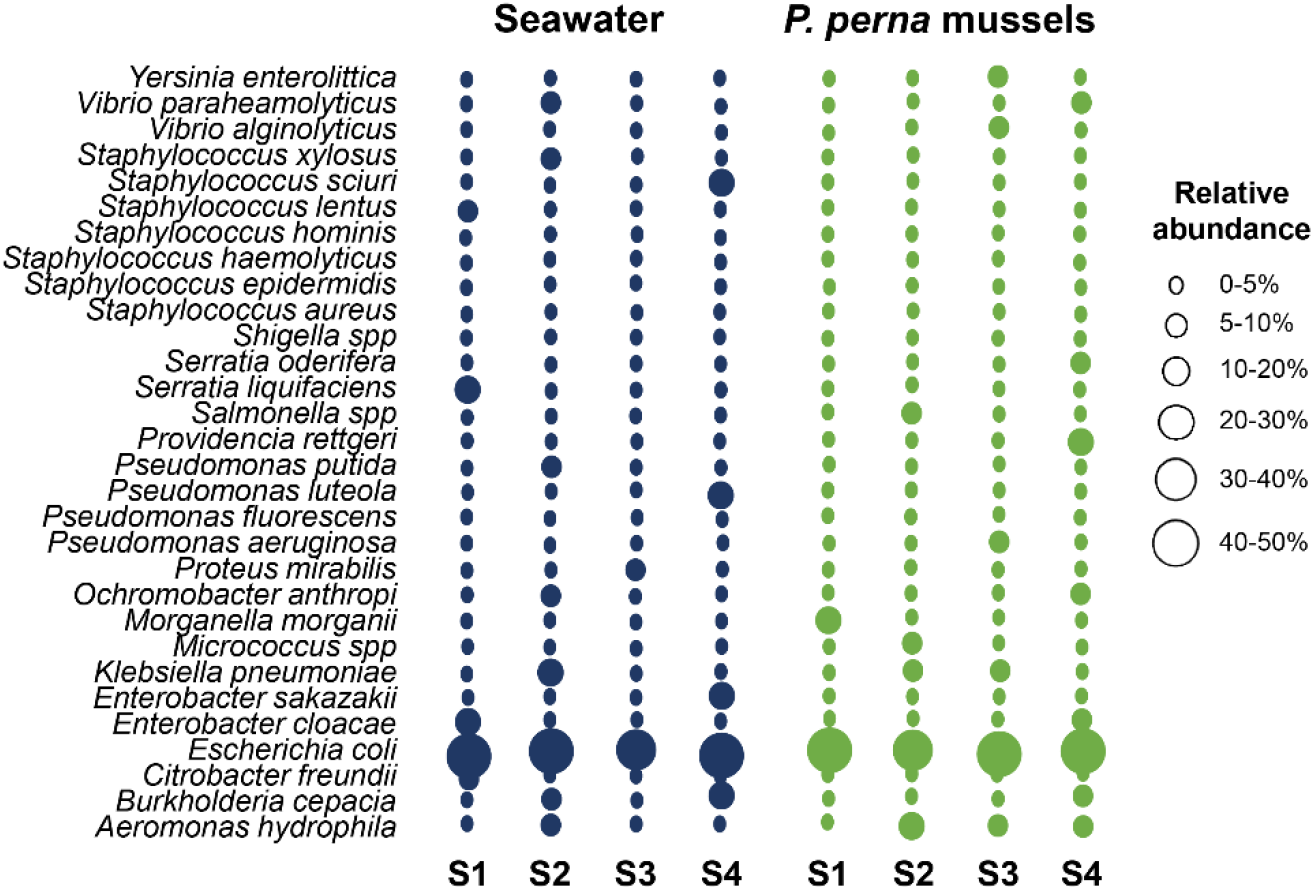
Relative abundances of potential pathogenic bacteria in seawater (blue) and *Perna perna* (green) samples. (S1) Cap de Garde (S2) Rezgui Rachid, (S3) Sidi Salem, (S4) Lahnaya.

### Molecular identification of selected isolates

In addition to biochemical identification 25 isolates of either Staphylococci or γ-proteobacteria were chosen for further identification via their 16S rRNA genes. Universal primers 27F and 1492R were used to PCR-amplify the 16S rRNA genes, and the products of approximately 1500 bp (Fig. 5) were Sanger sequenced. The 16S rRNA sequences were compared to the NCBI database, using BLAST. All sequences had between 97 and 100% identity to known bacterial species, permitting the identification of the analysed strains (Table 2). A phylogenetic tree was generated to visualize the evolutionary placement of our environmental bacteria with respect to their closest studied relatives (Fig. 6). A main clade, with high bootstrap value (100% bootstrap) grouped 17 isolates of seven genera within the family of Enterobacteriaceae; namely *Escherichia/Shigella, Klebsiella, Enterobacter, Citrobacter, Proteus* and *Morganella*. The *Staphylococci* were only represented by two isolates (BM2 and BS4) which were close to type strain *S. epidermidis* ATCC 10145^T^ (100% bootstrap). Overall, most (18/25) of the 16S rRNA gene sequence identification results matched with the genus identification using API tests (Table 2).

**Fig. 5.**
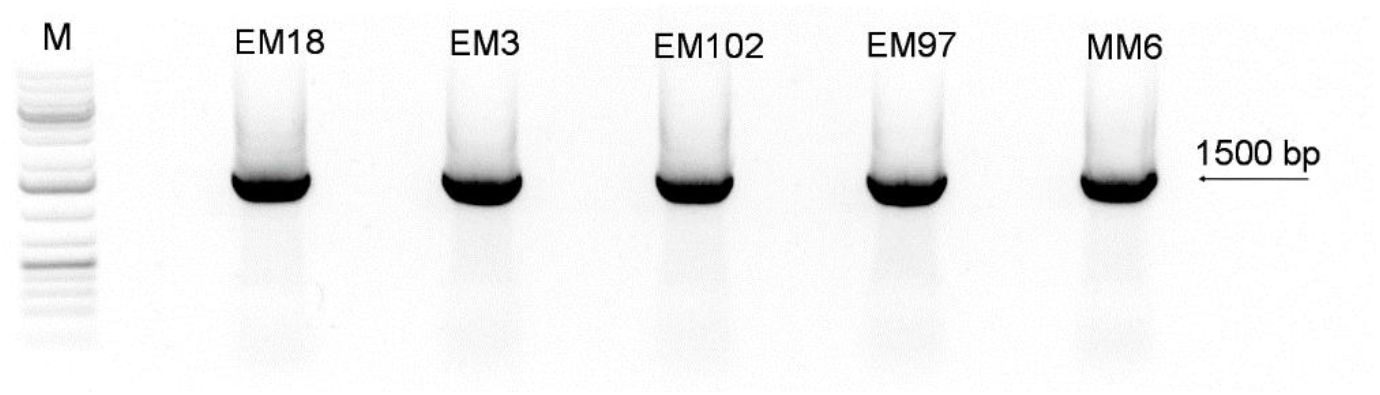
PCR amplification of the 16S rRNA gene. M: DNA ladder, lanes (EM18-MM6) represent amplified product (approx. 1500bp) of *E. coli* isolates.

**Fig. 6.**
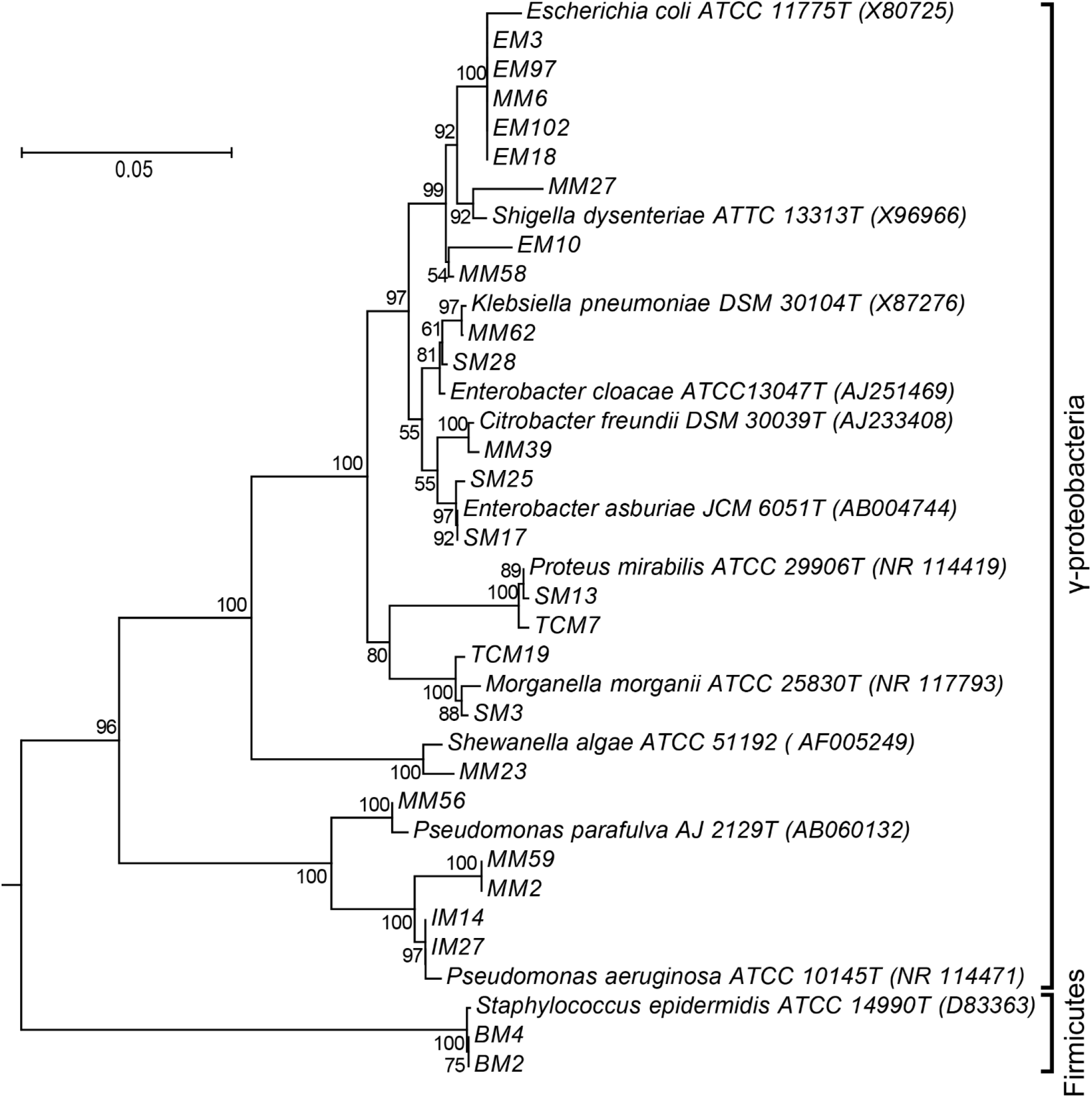
Phylogeny of the 25 isolates with molecular identification. 16S rRNA sequences from the 25 selected isolates together with the best hit from the GenBank database for each of the sequences, were compared in a Clustal X multiple sequence alignment (Kumar et al. 2016). Accession numbers of the reference sequences are in parentheses and Halobacterium sp. A1T was used to root the tree.

### Multi Locus Sequence Typing analysis (MLST)

*E. coli* comprised more than 40% of the isolated strains, and several individual *E. coli* isolates came from the very same environmental context (i.e. same sampling-site, sample-date, and environmental compartment). We therefore wondered whether these *E. coli* isolates were due to multiple separate contamination events or were caused by a single highly abundant *E. coli* strain which was able to thrive and outcompete other bacteria in the given condition. To test whether the five isolates (EM3, EM18, EM97, EM102, and MM6) isolated from *P. perna* mussels at Site 3 may belong to the same strain of *E. coli*, we PCR-amplified (Fig. 7) and Sanger-sequenced sections of seven conserved genes (*trpA, trpB, dinB, polB, putP, pabB and icdA*). A tree based on a multiple alignment of the concatenated sequences from our five strains (and the equivalent gene-sections from other *E. coli* strains) revealed that our isolates were most closely related to each other, and slightly more distantly to *E. coli* strains K12 and SCU-103 (Fig. 8).

**Fig. 7.**
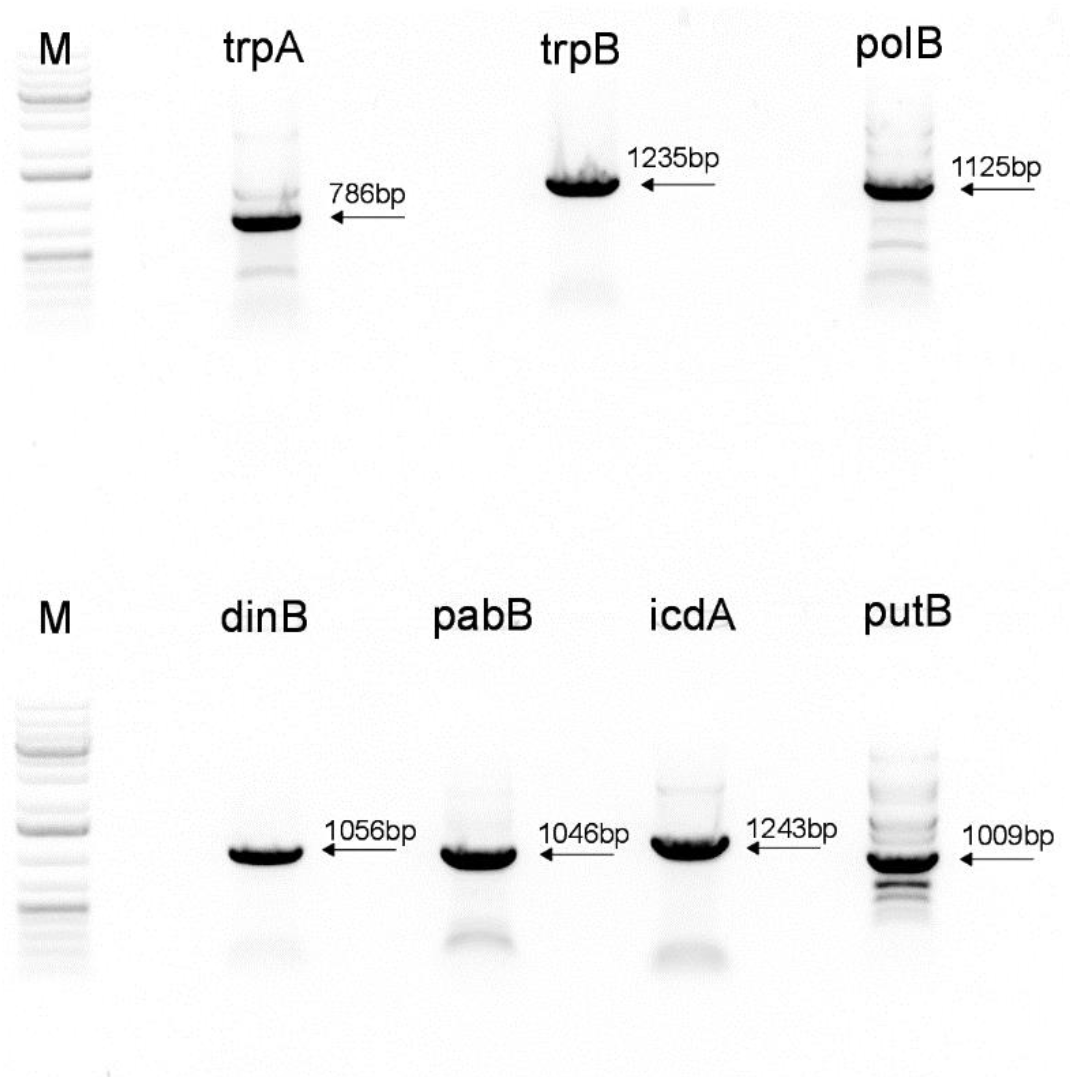
PCR amplification of seven genes of *E. coli* EM97. Lanes *trpA* to *putB* represent PCR products amplified from the EM97 isolate (*E. coli*). M: DNA ladder.

**Fig. 8.**
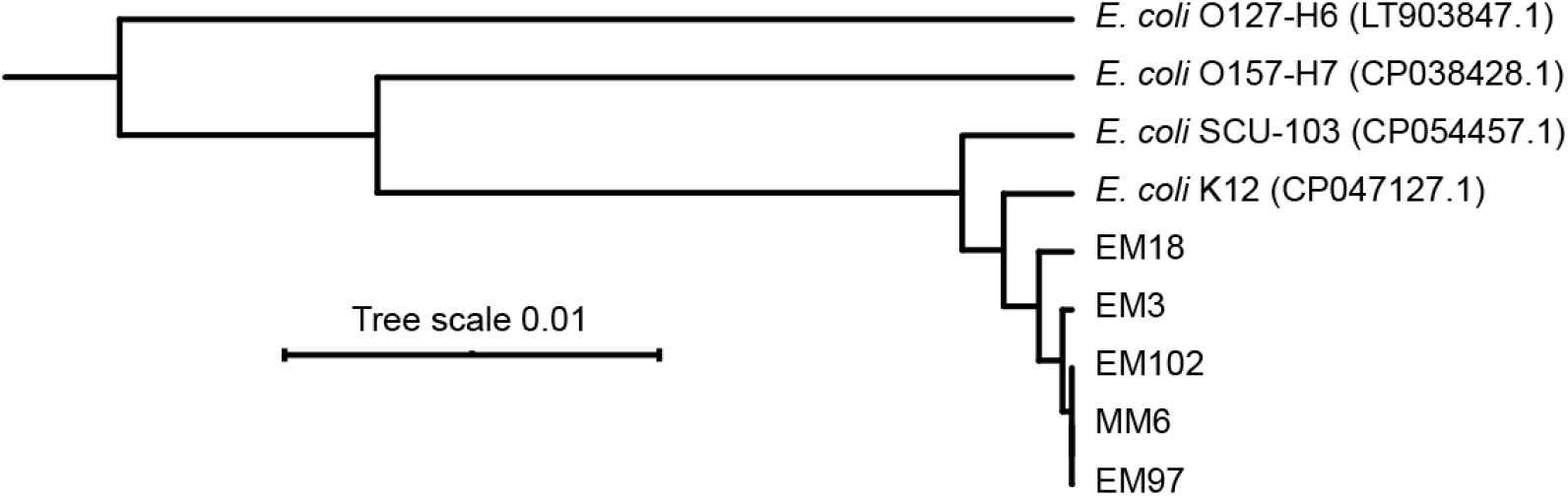
The *E. coli* strains isolated from S3 are similar but not identical. Phylogenetic tree showing the distances between the five analysed *E. coli* strains from S3, compared to *E. coli* strains from the NCBI database (accession numbers in parentheses). The tree was rooted using *Salmonella enterica*.

### Statistical analysis

The results of Spearman’s correlation analysis between FIB and physicochemical variables are given in Table 3. According to the correlation coefficients, FS appeared to be the most correlated indicator with all environmental variables except pH and SS: DO (r=-0.72, p<0.0001); temperature (r=0.64, p<0.0001) and salinity (r=0.46, p<0.01). In contrast, EC and TC were found to be positively and significantly correlated with SS (r=0.51 and 0.57; respectively, and both with p<0.001) and negatively correlated with DO (r=-0.46, p<0.01; r=-0.37, p<0.01; respectively). Principal Component Analysis (PCA) results revealed that the three first main components together explain 92.4% of the total information (Fig. 9a and b). The first PC, which represents 56.3% of the variance, was the most significant component of the latter. It was mainly loaded by the Temperature (0.95), Salinity (0.89), Dissolved Oxygen (−0.77), SS (−0.66) and FS (0.55). The second PC, representing 22.4% of the variation, was found to be positively correlated by pH (r = 0.86). The third PC, representing 13.7%, was positively correlated with TC and EC (r= 0.61 and r=0.52, respectively). According to the PCA plot a clear opposition was observed between the 12 months of sampling and the distribution of the four sites on the first two axes. S2 and S3 were strongly correlated with each other and showed maximum variations of fecal contamination during the warm months of the year, whereas S1 and S4 demonstrated lower fecal contamination variations during the cold months (Fig. S1).

**Fig. 9.**
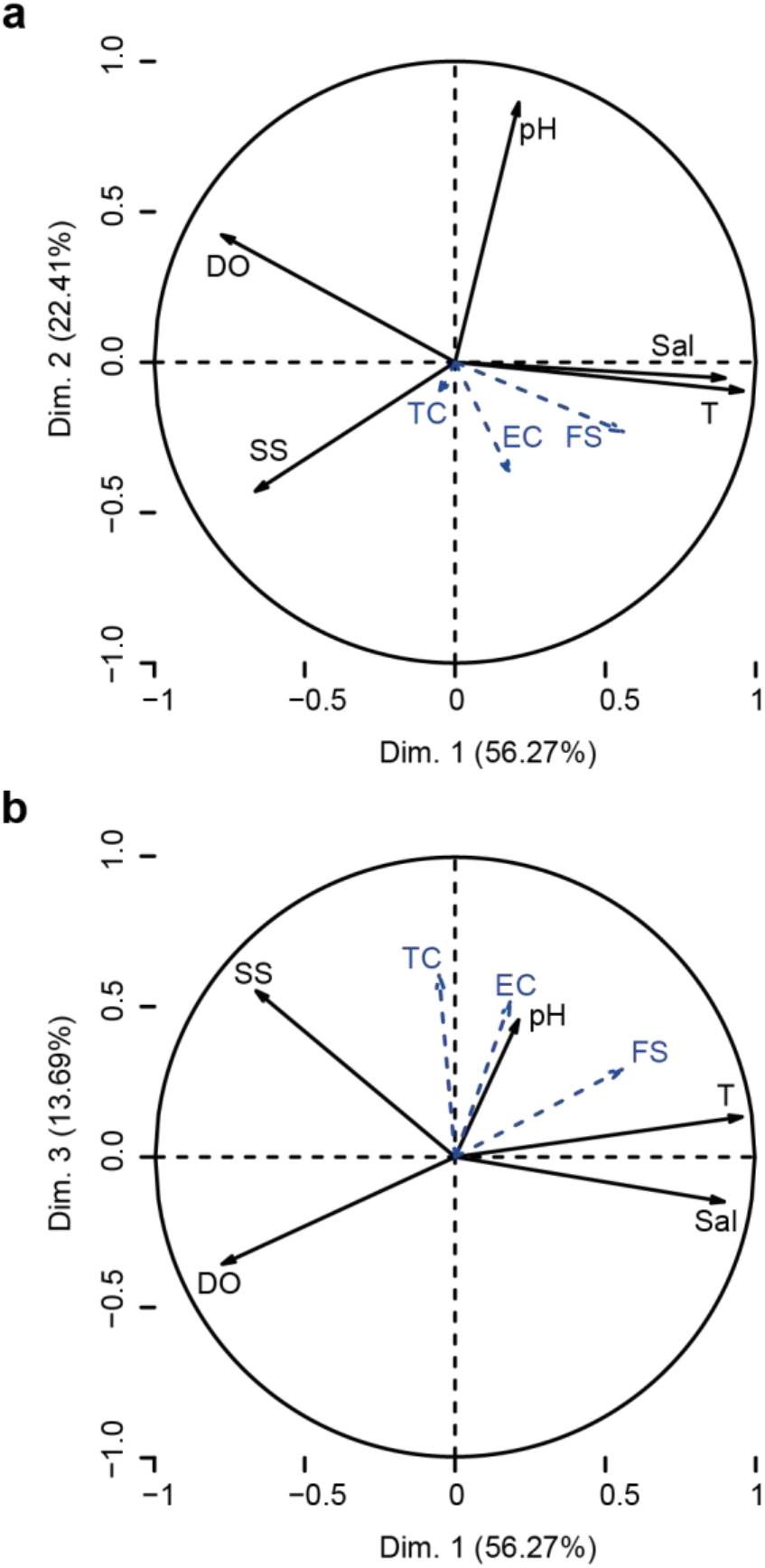
Principal component analysis (PCA) performed on data from seawater samples. **a** Correlation of environmental variables with the two first axes of the standard PCA. **b** Correlation of environmental variables with the first and third axes of the standard PCA. The main variables are indicated with solid arrows and supplemental variables are indicated with dotted arrows. DO: dissolved oxygen, Sal: salinity, T: water temperature, SS: Suspended solids. EC: *Escherichia coli*, TC: total coliforms and FS: fecal streptococci.

## Discussion

The results of this study revealed a significant increase of fecal contamination in the Gulf of Annaba compared to previous studies conducted in the same area (Hidouci et al. 2014, Kadri et al. 2015, 2017) and in other coastal regions of the Mediterranean Sea (Bouhayene et al, 2014; Boutaib et al. 2015; Dallarés al. 2018; Rincé et al, 2018). Much of this difference is probably due to the continuous pollution pressure in the Gulf, mainly related to anthropogenic activities, as well as rapid urbanization over the last few years. According to Grimes (2003) and Inal et al. (2018), nearly 19 million people (45% of the Algerian population) is living along the Gulfs near the largest agglomerations such as Oran, Algiers, and Annaba. In the current study, these impacts were differently manifested depending on the local sources of pollution at each site. The strong presence of FIB at S3 could be explained by significant urban, industrial and agricultural discharges that Wadi Seybouse (the second longest Wadi in Algeria) drains from its catchment basin of about 6470 km^2^ (ABH-CSM 1999; Mebarki 2000). Besides, higher bacterial concentrations may also be attributed to untreated wastewater effluents from a large part of Annaba city and its outskirts (Bouhamra, Joannoville and, its slaughterhouse) discharged directly into the sea via Wadi Bedjima, and also to the presence of a large colony of seabirds and animals (Telailia 2014). A similar increase in fecal contamination was reported in the same area by Kadri and coworkers (2017). Our present results further demonstrated that the overall contamination was exceptionally high at S2, revealing that the waters of this site should be considered as unsafe for bathing in accordance with the Algerian Bathing Water executive decrees (JORA 1993, 2006). Similar to S3, these high levels of FIB were primarily due to the domestic wastewaters from nearby homes discharged directly into the sea without prior treatment (Kadri et al. 2017). On the other hand, the lower concentrations observed in S1 and especially S4 are due to their remoteness from the city area and their particular hydrodynamics which may contribute to the dispersion of fecal pollutants in the water column (Kadri et al. 2015). Nevertheless, it is important to note that these two sites are often visited by swimmers in summer which explains the increasing levels of FIB during this period of the year (Kadri et al. 2017). According to several studies, fecal contamination at bathing beaches can be hazardous to humans because many pathogenic bacteria could be ingested during recreational water activities leading to various waterborne diseases (Marion et al. 2010; Santhiya et al. 2011; Arnold et al. 2016). It is, therefore, eminent to take exceptional protective measures by the government (proposing specific treatment solutions, especially for wastewater discharges, prohibiting direct industrial and agricultural discharges, and improving public awareness) so that these areas have optimal quality levels to avoid any health risks. Data obtained showed that the levels of FIB were alarmingly higher in *P. perna* than in the surrounding seawaters during all the study period. These outcomes are consistent with those reported in other coastal regions worldwide, suggesting that this strong accumulation capacity is mainly related to the filter-feeding behavior of these sentinel organisms, which make them one of the best bio-indicators of fecal pollution in coastal waters (Stabili et al. 2005; Martinez and Oliveira 2010; Jayme et al. 2016; Bozcal and Dagdeviren 2020). Furthermore, the levels of intestinal indicators in all sampling sites were well above the permissible limits recommended according to Regulation (854/2004/EC) of 29 April 2014 for human consumption, which recommends less than 230 *E. coli*/100g. Thus, the mussels inhabiting the Gulf of Annaba would be unfit for direct consumption. FS were present throughout the entire sampling period with concentrations higher than *E. coli* in almost all samples, and this is also in agreement with previous reports (Tiefenthaler et al. 2009; Zegmout et al. 2011; Kadri et al. 2017; Islam et al. 2017). These germs are known to have a better survival period than fecal coliforms in surface waters as well as in the digestive tract of bivalves (Geldreich 1976, Noble et al. 2004). In addition to anthropogenic activities, the survival was also influenced by a multitude of natural variables, among them; the climatic changes was very likely responsible for the observed differences in bacterial concentrations in our samples. Indeed, our results demonstrated that elevated temperatures in the last spring and summer were associated with maximum FIB rates in both compartments. According to the Intergovernmental Panel on Climate Change (IPCC) Fifth Assessment Report (2013), Algeria will experience an increase in temperatures between 1 and 3.7 °C over the next few years. Consequently, this may lead to enhance the proliferation and persistence of bacteria for a long time in seawater and cause the deterioration of water resources (Mohammed and Al-Amin 2018, Barreras et al. 2019). This hypothesis was confirmed in the current study by significant and positive correlations between *E. coli* and FS, and the temperature revealed by Spearman correlation test (p < 0.05) and PCA analysis, which is consistent with other studies (Koirala et al. 2008; Gutiérrez-Cacciabue et al. 2014, Abia et al. 2015; Islam et al. 2017). Another probable reason for the high levels of FIB in *P. perna* in this period is probably the physiology of this sentinel species. Burge et al. (2016) have indicated that elevated temperatures promote the filtration rates in mussels and, therefore, they can retain more microorganisms from the surrounding waters. Besides the temperature, salinity is also a crucial variable for the survival of FIB in aquatic environments. In our study, only FS showed a significant positive correlation (p < 0.001) with salinity (Table 3). These germs are known for their high resistance to harsh environmental stressors and tolerance to high concentrations to salt, making them powerful indicators of fecal contamination (Byappanahalli et al. 2012). Conversely, DO show the strongest correlations with all groups of FIB, mainly due to the bacterial degradation of detritus which consumes a lot of oxygen. This biodegradation was more important with the increase in temperature in summer (5 mg/L) (Fig. 2), especially in highly contaminated sites, which receive massive quantities of domestic discharges and industrial effluents. These findings are in agreement with those of a recent study by Chávez-Díaz et al. (2020), which found negative correlations between DO and FIB. The latter were found to be positively correlated with SS, which, according to the literature, play a protective role for intestinal bacteria against solar radiation and predators (Walters et al. 2014; Kadri et al. 2017). This appears to be the case for the SS-rich waters of S2 and S3, which reportedly contain large quantities of FIB. Numerous studies have indicated that FIB are used as surrogates to estimate the possible presence of pathogenic microorganisms, especially when they are found at high levels (Wilkes et al. 2011; Shoults and Ashbolt 2018). In Algeria and especially in the Gulf of Annaba, very few studies on the presence of bacterial pathogens in the mussels and recreational waters, and their human health risks were investigated. Similar to the study conducted by Stabili et al. (2005) and Cavallo et al. (2008) in the Northern Ionian Sea of Italy, the bacterial community of *P. perna* mussels from all sites in our study was very similar to that of surrounding waters but with higher abundance. Proteobacteria was the most dominant phylum (88/208, 46%) represented mainly by members of the family *Enterobacteriaceae* and divided into two major groups: enteric and marine or environmental bacteria. Among enteric bacteria, *Salmonella*, *Shigella* and, *E. coli* are known to be the bacteria that are most involved in human gastrointestinal tract infections. According to Yang et al. (2017), 1.7 billion cases of human diarrhea caused primarily by pathogenic strains of *E.coli* have been recorded worldwide each year. Similarly, Sánchez-Vargas et al. (2011) and Neogi et al. (2014) reported that *Salmonella typhi* causing enteric fever affected approximately 450 per 100,000 children in India and Pakistan. These results indicated that domestic wastes, especially from the most polluted sites are most likely the primary source of pollution in the Gulf of Annaba since enteric bacteria are mainly derived from the excrement of warm-blooded animals, including humans (Poharkar et al. 2017). Environmental bacteria such as *Pseudomonas*, *Aeromonas*, and *Shewanella* were also identified in both environmental compartments. Species of the genus *Aeromonas* are widely isolated from aquatic environments and frequently reported to cause waterborne and seafood infections (gastroenteritis and septicemia) (Chopra and Houston 1999; Joseph et al. 2013; Hamid et al. 2016). *Pseudomonas* spp. are another ubiquitous microorganisms found in marine shellfish and recreational waters (Maravić et al. 2018; Goh et al. 2019). These multidrug-resistant pathogens have been previously reported to be associated with diarrhea, intra-abdominal and nosocomial infections, particularly in immune-compromised patients (Morrissey et al. 2013; Streeter and Katouli 2016). The results of biochemical identification also revealed the detection of different species of the genus Vibrio in the four sampling sites, of which *V. paraheamolyticus* was the most isolated microorganism. Our findings are consistent with the results of numerous studies conducted worldwide (Stabili et al. 2005; Esteves et al. 2015; Vezzulli et al. 2018; Nguyen et al. 2018; Hackbusch et al. 2020). *Vibrio*. spp are waterborne bacteria naturally found in estuarine and coastal environments. Yet, certain species can be pathogenic to humans and marine organisms such as bivalves (Eggermont et al. 2017; Rincé et al. 2018; Bozcal and Dagdeviren, 2020). In August 2018, the Algerian Ministry of Health reported a cholera outbreak in Blida and five other regions (Algiers, Tipaza, Bouira, Médéa, and Ain Defla) in the north of the country. This devastating and strictly human epidemic caused mainly by *V. cholera* O1 or O139 can cause serious outbreaks such as severe dehydrating diarrhea and even death (Feldhusen 2000). In the Gulf of Annaba, this germ was found in S3 mussels in the same period of the outbreak classifying this site as the area of highest risk. In addition to *Enterobacteriaceae*, 24 isolates of the genus *Staphyloccocus* were also detected during the study period. These germs, including *S. aureus*, are well-known causative agents of several diseases in humans such as skin rashes, pneumonia, ear and eye infections, endocarditis and, meningitis (Schets et al. 2020; Yaghoubzadeh et al. 2020). According to Pomykala et al. (2012), some coagulase-positive staphylococci are common seafood pathogens and may pose a significant risk to human health through improper consumption of bivalve mollusks. Furthermore, methicillin-resistant *S. aureus* (MRSA), which is one of the most harmful pathogens on human health, has also been frequently detected on several recreational beaches in the United States (Abdelzaher et al. 2010; Levin-Edens et al. 2012; Plano et al. 2013; Thapaliya et al. 2017). As mentioned above, the biochemical identification results revealed that isolated bacteria were primary members of the *Enterobacteriaceae* family. Strains of this large group of bacteria are known to be closely related to each other and difficult to distinguish by conventional methods (Nhung et al. 2007; Hamdi et al. 2017). Besides, the use of biochemical identification alone can be problematic as some new taxa may not be included in available databases (Janda and Abbott 2002). For this reason, additional molecular identification targeting the 16S rRNA gene was performed on 25 strains, including two Gram-positive bacteria isolated from *P. perna* mussels of S3. In general, the biochemical identification at the genus level was confirmed in 72% of the cases by the 16S rRNA gene sequencing (Table 2). This molecular method proved to be more accurate for bacterial identification as all strains exhibited more than 97% sequence similarities with their matching sequences retrieved from the GenBank database. The ribosomal 16S rRNA gene has highly conserved regions in all bacterial cells, interspersed with nine hyper-variable stretches of sequences (V1-V9), and is a molecular fingerprint for bacterial identification and taxonomic classification (Benga et al. 2014; Jo et al. 2016; Monticelli et al. 2019). This method also has minor limitations such as high sequence similarities among closely related species (Jo et al. 2016). Therefore, the use of more than one target gene, especially genes that are more susceptible to genomic drift than the 16S rRNA gene, provide a more detailed differentiation between closely related bacterial isolates. For the strains (EM3, EM18, EM97, EM102, and MM6) identified as *E. coli* using API tests and 16S rRNA gene sequencing, MLST was performed to further understand their phylogenetic relationships. Interestingly, the results indicate that our isolates were very similar to each other but nevertheless distinct, and and therefore did not belong to the same strain of *E. coli* (Fig. 8). This suggests that they came from a variety of separate human and animal sources of fecal contamination, since *E. coli* is mainly found in the fecal wastes of warm-blooded mammals (Poharkar et al. 2017). Therefore, the use of Microbial Source Tracking (MST) technique to identify both human and animal specific markers in future studies will be an important tool for understanding the origin of fecal pollution in the Gulf of Annaba, and for assessing the associated health risks related to the presence of pathogenic microorganisms.

## Conclusion

The current study revealed that the presence of fecal indicators in the marine waters and mussels of the Gulf of Annaba was strongly affected by both anthropogenic activities and environmental variables. Multiple analysis showed that *P. perna* was the most contaminated sample with the highest levels of FIB in all sampling sites, especially those located inside the Gulf (S2 and S3) near Annaba. These principal findings validate our choice to use this species as an effective bio-indicator to assess the microbial quality of coastal waters. Our results also demonstrated that different pathogenic bacteria were detected during the study period. The survival and the presence of these infectious agents in *P. perna* is a matter of great concern regarding epidemic diseases that they may occur when consumed by humans. Therefore, the implementation of necessary measures should be carried out, especially in highly polluted sites in order to protect environmental resources and human health.

## Declarations

### Ethics approval and consent to participate

Not applicable

### Consent for publication

Not applicable

### Availability of data and materials

The datasets used and/or analysed during the current study are available from the corresponding author (Mouna Boufafa) on reasonable request.

### Authors’ contributions

MBo performed the experiments. MBo, SK and MBe analysed the environmental sampling data. PR analysed the MLST data. MBo, SK, PR and MBe wrote the paper.

## Acknowledgements

The authors would like to thank Dr. Hassen Touati and Dr. Fatma Guellati Bouyema for their assistance in the statistical analysis of data.

---

**Table 1.**
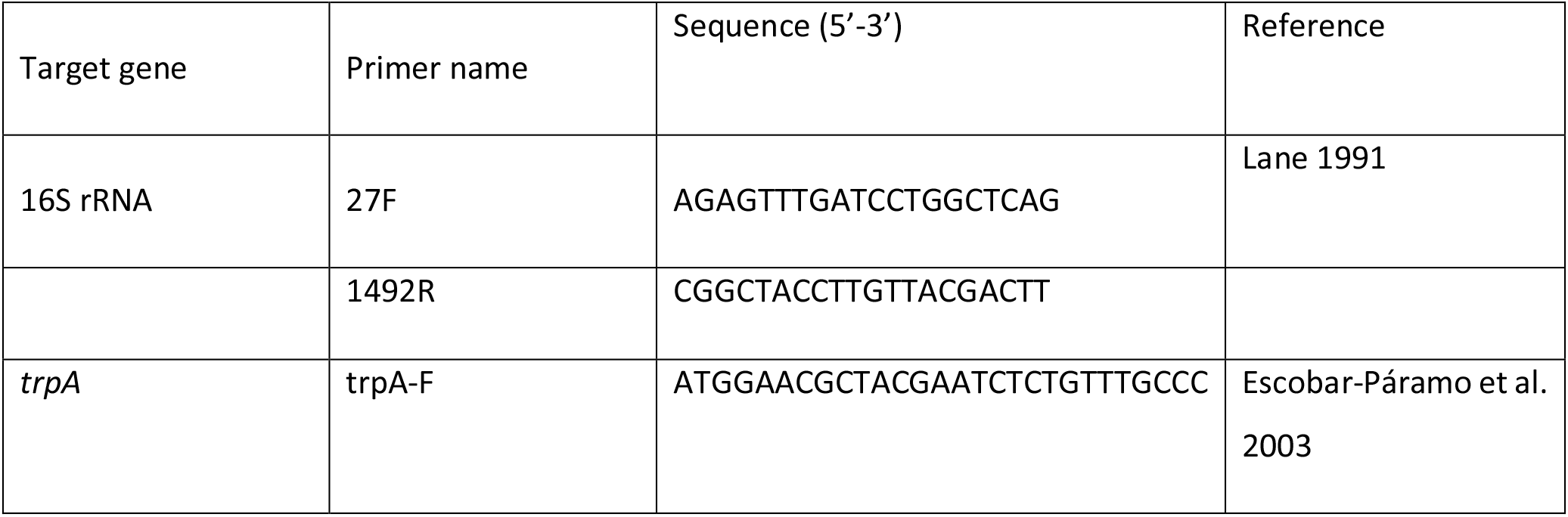
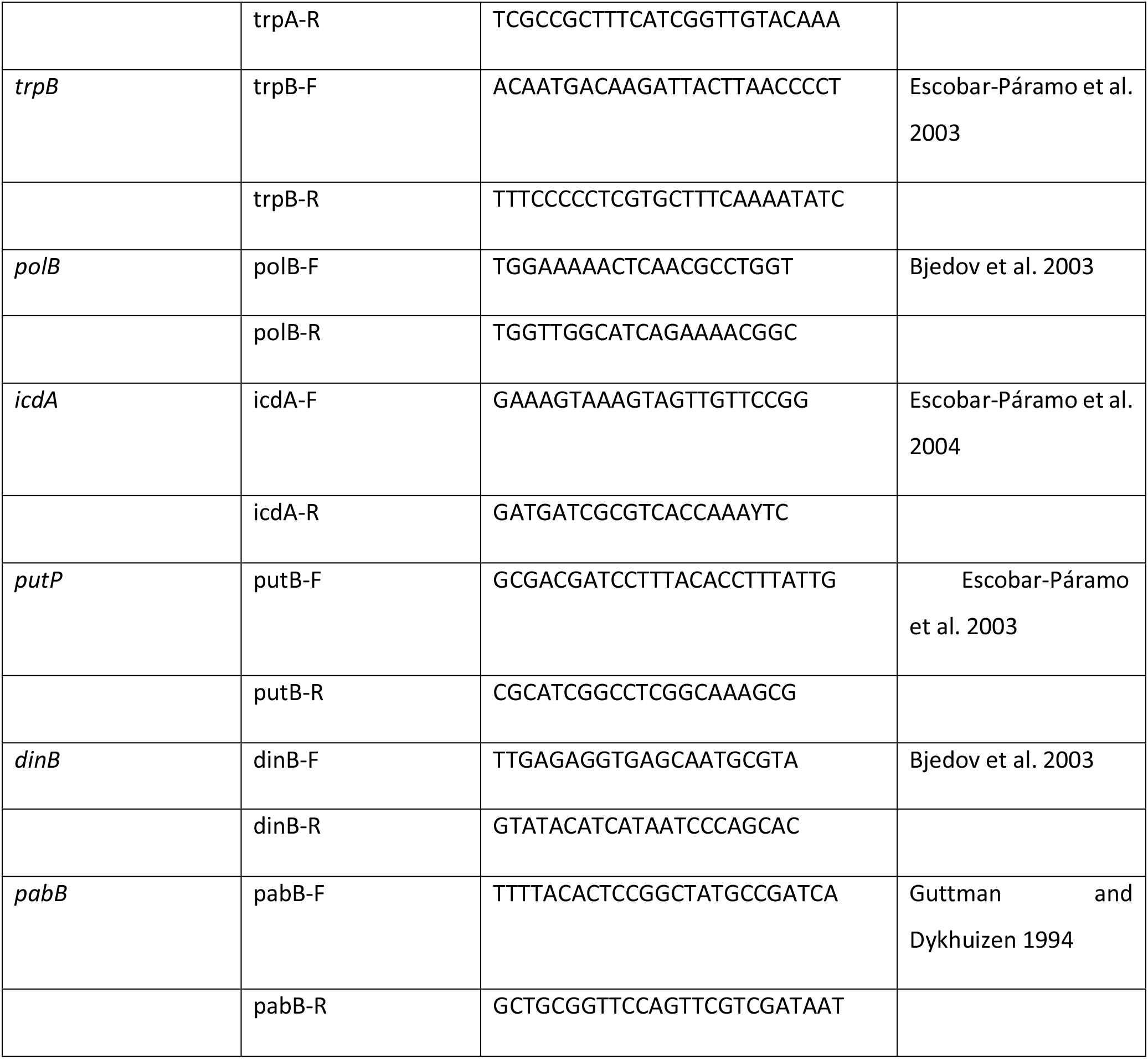
Primers used in this study

**Table 2.** Biochemical and molecular identification of 25 isolates isolated from *P. perna* mussels

| Isolate code | Api identification | Best hit to 16S rRNA sequence | MLST identification |
| --- | --- | --- | --- |
| EM3 | *Escherichia coli* | *Escherichia coli* | *Escherichia coli* |
| MM6 | *Escherichia coli* | *Escherichia coli* | *Escherichia coli* |
| EM97 | *Escherichia coli* | *Escherichia coli* | *Escherichia coli* |
| EM102 | *Escherichia coli* | *Escherichia coli* | *Escherichia coli* |
| EM18 | *Escherichia coli* | *Escherichia coli* | *Escherichia coli* |
| EM10 | *Escherichia coli* | *Escherichia coli* | NA |
| MM58 | *Escherichia coli* | *Escherichia coli* | NA |
| TCM7 | *Aeromonas hydrophila* | *Proteus mirabilis* | NA |
| MM59 | *Chromobacterium violaceum* | *Pseudomonas aeruginosa* | NA |
| SM25 | *Citrobacter koseri* | *Enterobacter asburiae* | NA |
| SM28 | *Enterobacter cloacea* | *Enterobacter cloacea* | NA |
| MM62 | *Klebsiella pneumoniae* | *Klebsiella pneumoniae* | NA |
| SM3 | *Morganella morganii* | *Morganella morganii* | NA |
| TCM19 | *Ochromobacter anthropi* | *Morganella morganii* | NA |
| IM14 | *Pseudomonas aeruginosa* | *Pseudomonas aeruginosa* | NA |
| MM56 | *Pseudomonas fluorescens* | *Pseudomonas parafulva* | NA |
| IM27 | *Pseudomonas luteola* | *Pseudomonas aeruginosa* | NA |
| MM2 | *Pseudomonas putida* | *Pseudomonas aeruginosa* | NA |
| SM13 | *Proteus mirabilis* | *Proteus mirabilis* | NA |
| MM39 | *Salmonella choleraesuis* | *Citrobacter freundii* | NA |
| MM23 | *Salmonella spp* | *Shewanella algae* | NA |
| SM17 | *Serratia odorifera* | *Enterobacter asburiae* | NA |
| MM27 | *Shigella spp* | *Shigella dysenteriae* | NA |
| BM2 | *Staphylococcus lentus* | *Staphylococcus epidermidis* | NA |
| BM4 | *Staphylococcus epidermidis* | *Staphylococcus epidermidis* | NA |

NA: Not analysed

**Table 3.** Spearman’s correlation matrix of the seawater quality variables in 2018

|  | Temperature | Salinity | pH | Dissolved Oxygen | Suspended Solids | Total Coliforms (TC) | Escherichia coli (EC) |
| --- | --- | --- | --- | --- | --- | --- | --- |
| Salinity | 0.824*** |  |  |  |  |  |  |
| pH | 0.181 | 0.185 |  |  |  |  |  |
| Dissolved Oxygen | -0.836*** | -0.650*** | 0.008 |  |  |  |  |
| Suspended Solids | -0.483*** | -0.529*** | -0.223 | 0.183 |  |  |  |
| Total coliforms (TC) | 0.137 | -0.021 | 0.183 | -0.371** | 0.565*** |  |  |
| Escherichia coli (EC) | 0.285* | 0.140 | 0.025 | -0.459** | 0.507*** | 0.769*** |  |
| Fecal Streptococci (FS) | 0.643*** | 0.461*** | 0.119 | -0.721*** | -0.025 | 0.559*** | 0.453*** |

\* $p \leq 0.05$ ; \*\* $p \leq 0.01$ ; \*\*\* $p \leq 0.001$ .

## SUPPLEMENTARY MATERIAL

**Figure S1.**
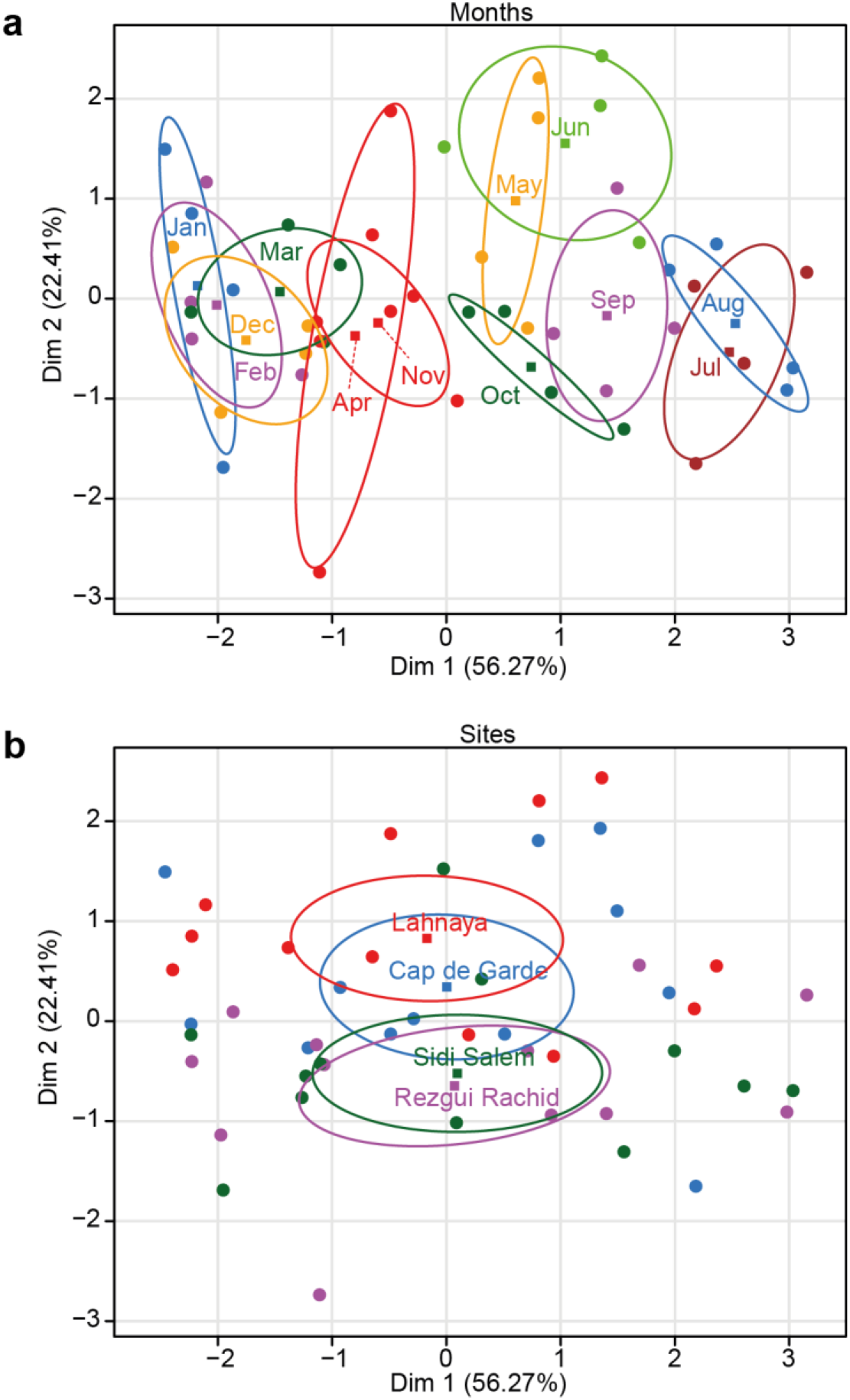
Sampling month and site projections. **a** Sampling month projection on the two first axes of the standard PCA. **b** Sampling site projection on the two first axes of the standard PCA.

**Table S1.** Identification results of pathogenic bacteria in seawater and *Perna perna* mussels

| Species |  |  | S1 | S1 | S2 | S2 | S3 | S3 | S4 | S4 | Total |
| --- | --- | --- | --- | --- | --- | --- | --- | --- | --- | --- | --- |
|  |  |  | W | M | W | M | W | M | W | M | W/M |
| *Aeromonas hydrophila* |  |  |  | 1 | 1 | 5 | 1 | 3 |  | 1 | 12 |
| *Burkholderia cepacia* |  |  |  |  | 1 |  | 1 | 1 | 1 | 1 | 5 |
| *Buttiauxella agresis* |  |  | 1 |  |  |  |  |  |  |  | 1 |
| *Citrobacter amalonaticus* |  |  |  |  |  |  |  | 1 |  |  | 1 |
| *Citrobacter freundii* |  |  | 1 |  |  |  |  | 1 |  |  | 2 |
| *Enterobacter cloacae* |  |  | 2 | 1 |  |  | 1 | 1 |  | 1 | 6 |
| *Enterobacter sakazaki* |  |  |  |  |  | 1 |  |  | 1 |  | 2 |
| *Escherichia coli* |  |  | 6 | 11 | 8 | 13 | 8 | 28 | 4 | 8 | 86 |
| *Klebsiella pneumoniae* |  |  |  |  | 2 | 3 |  | 3 |  |  | 8 |
| *Klebsiella oxytoca* |  |  |  |  |  |  |  |  | 1 |  | 1 |
| *Kluyvera spp* |  |  |  | 1 |  |  |  |  |  |  | 1 |
| *Micrococcus spp* |  |  |  |  |  | 3 | 1 |  |  |  | 4 |
| *Morganella morganii* |  |  |  | 4 |  |  |  | 1 |  |  | 5 |
| *Ochromobacter anthropi* |  |  |  |  | 1 |  |  |  |  | 1 | 2 |
| *Pasteurella pneumotropica* |  |  |  |  |  |  | 1 |  |  |  | 1 |
| *Pantoea spp2* |  |  |  |  |  |  | 1 |  |  |  | 1 |
| *Proteus mirabilis* |  |  |  |  |  |  | 2 | 1 |  |  | 3 |
| *Proteus vulgaris* |  |  |  |  |  | 1 |  | 1 |  |  | 2 |
| *Providencia rettgeri* |  |  |  |  |  |  |  |  |  | 2 | 2 |
| *Pseudomonas aeruginosa* |  |  |  | 1 |  | 1 | 1 | 4 |  |  | 7 |
| *Pseudomonas fluorescens* |  |  |  |  |  | 1 | 1 |  |  |  | 2 |
| *Pseudomonas luteola* |  |  |  | 1 |  | 2 |  |  | 1 |  | 4 |
| *Pseudomonas putida* |  |  |  |  | 1 |  |  | 1 |  |  | 2 |
| *Salmonella choleraesuis spp arizonae* |  |  |  |  |  |  |  | 1 |  |  | 1 |
| *Salmonella spp* |  |  |  |  |  | 3 |  |  |  |  | 3 |
| *Serratia liquefaciens* |  |  | 2 |  |  |  |  |  |  |  | 2 |
| *Serratia marcesens* |  |  |  |  |  |  | 1 |  |  |  | 1 |
| *Serratia odorifera* |  |  |  |  |  |  |  | 1 |  | 1 | 2 |
| *Serratia plymuthica* |  |  | 1 |  |  |  |  |  |  |  | 1 |
| *Shigella spp* |  |  |  | 1 |  | 2 |  |  |  |  | 3 |
| *Staphylococcus aureus* |  |  |  |  |  | 1 | 1 |  |  |  | 2 |
| *Staphylococcus capitis* |  |  |  |  |  | 1 |  |  |  |  | 1 |
| *Staphylococcus epidermidis* |  |  |  |  |  | 2 |  |  |  |  | 2 |
| *Staphylococcus haemolyticus* |  |  |  | 1 |  |  | 1 |  |  |  | 2 |
| *Staphylococcus hominis* |  |  |  |  |  |  | 1 | 1 |  |  | 2 |
| *Staphylococcus lentus* |  |  | 1 |  |  |  |  | 1 |  |  | 2 |
| *Staphylococcus lococcus* |  |  |  |  |  | 1 |  |  |  |  | 1 |
| *Staphylococcus sciuri* |  |  |  |  |  |  | 1 | 2 | 1 |  | 4 |
| *Staphylococcus xylosus* |  |  |  |  | 1 | 1 | 1 | 1 |  |  | 4 |
| *Staphylococcus warneri* |  |  |  |  |  | 1 |  |  |  |  | 1 |
| *Vibrio alginolyticus* |  |  |  |  |  |  | 2 |  |  |  | 2 |
| *Vibrio cholerae* |  |  |  |  |  |  |  | 1 |  |  | 1 |
| *Vibrio metschnikovii* |  |  |  | 1 |  |  |  |  |  |  | 1 |
| *Vibrio parahaemolyticus* |  |  |  |  | 1 | 1 | 1 | 2 |  | 1 | 6 |
| *Vibrio vulnificus* |  |  |  | 1 |  |  |  |  |  |  | 1 |
| *Yersinia enterocolitica* |  |  |  |  |  |  |  | 3 |  |  | 3 |
| **Total** |  |  | **14** | **24** | **16** | **43** | **27** | **59** | **9** | **16** | **208** |
| Table 1 (supplementary materiel): identification results of pathogenic bacteria in seawater and Perna perna mussels) |  |  |  |  |  |  |  |  |  |  |  |
| W: Seawater | M: Mussels | Yellow background indicates the pathogenic bacteria from Figure 4 |  |  |  |  |  |  |  |  |  |

